# The effect of number of healthcare visits on study sample selection and prevalence estimates in electronic health record data

**DOI:** 10.1101/622761

**Authors:** Laura J. Rasmussen-Torvik, Al’ona Furmanchuk, Alexander J. Stoddard, Kristen I. Osinski, John R. Meurer, Nicholas Smith, Elizabeth Chrischilles, Bernard S. Black, Abel Kho

**Author notes:** Corresponding Author: Laura J. Rasmussen-Torvik, PhD, MPH, 680 N. Lake Shore Drive, Suite 1400, Chicago, IL 60611, p., f.

## Abstract

**Introduction:** Few studies have addressed how to select a study sample when using electronic health record (EHR) data.

**Methods:** Year 2016 EHR data from three health systems was used to examine how alternate definitions of the study sample, based on number of healthcare visits in one year, affected measures of disease period prevalence. Curated collections of ICD-9, ICD-10, and SNOMED codes were used to define three diseases.

**Results:** Across all health systems, increasing the minimum required number of visits to be included in the study sample monotonically increased crude period prevalence estimates. The rate at which prevalence estimates increased with number of visits varied across sites and across diseases.

**Conclusions:** When using EHR data authors must carefully describe how a study sample is identified and report outcomes for a range of sample definitions, so that others can assess the sensitivity of reported results to sample definition in EHR data.

## Introduction

Increased adoption of electronic health records (EHR) has generated increased interest in using these data in clinical and epidemiologic research ^1^. EHRs have been proposed to offer an efficient means for identifying eligible subjects for retrospective database studies and for prospective observational studies or pragmatic trials. EHRs offer data elements that are desirable for capturing baseline inclusion and exclusion criteria, covariates, treatments and interventions, and study outcomes.

EHRs can provide a reasonably complete picture of a patient’s health and medical encounters in “closed” health systems, such as traditional health maintenance organizations. However, many EHRs are drawn from non-closed systems, which likely provide only some of a patient’s health care encounters. The EHR will often reflect a subset (sometimes a small subset) of patient encounters and as a result, diagnoses. Therefore, it is challenging to define a study sample using EHR data, and, therefore, to calculate even basic outcomes such as chronic disease prevalence. Researchers must specify criteria for determining which patients have sufficient information in the EHR to be included in the study sample. Often, this sample is defined by requiring that persons have a minimum number of visits in a defined period (for example 2 visits in a 3 year period ^2^).

Many studies have been published that propose, validate, and, in some cases, compare disease definitions in EHR systems (see Sprat et al. ^3^ for a type 2 diabetes example and Pathak et al. for an overview ^4^). One study included a simulation demonstrating for lower-sensitivity phenotypes, the potential for bias is exacerbated when the medical condition also leads to more patient encounters^5^. However, the specific implications of different methods for selecting a study sample have received little examination. The objective of this report was to examine how changing criterion for number of visits in EHR data required for inclusion in a study sample would impact one basic epidemiologic measure: estimates of disease period prevalence, i.e. the proportion of individuals in a defined population that have a disease during a specified time period. Period prevalence of three common diseases was examined across three large geographically and demographically diverse health care systems.

## Methods

EHR data from three different health systems participating in the Greater Plains Collaborative ^6^ and Chicago-Area (CAPriCORN) ^7^ Clinical Data Research Networks (CDRNs) was used. Although the specific health systems are not identified here, systems with varying locations and diverse population demographic populations served were intentionally chosen. Each health system provided inpatient, outpatient and emergency department diagnoses (using ICD9, ICD10 and SNOMED codes) for all patients with health care encounters on two or more discrete days during 2016. The initial sample was filtered to require at least two “visits” (defined as any Ambulatory Visit, Emergency Department Visit, Emergency Department Admit to Inpatient Hospital Stay, Inpatient Hospital Stay, Non-Acute Institutional Stay, Observation Stay, Institutional Professional Consult, or Other Ambulatory visit, per the People-Centered Outcomes Research Institute Common Data Model (https://pcornet.org/pcornet-common-data-model/), and varied the minimum number of visits from 2 to 6.

To define cases of specified diseases in the study samples, curated collections of ICD-9, ICD-10, and SNOMED codes drawn from the Center for Medicare Studies Electronic Clinical Quality Measures (eCQMs—https://ecqi.healthit.gov/ecqms) were used. For myocardial infarction, codes for the denominator of the eQCM “Coronary Artery Disease: Beta-Blocker Therapy-Prior Myocardial Infarction (MI) or Left Ventricular Systolic Dysfunction (LVEF <40%)” were used; for diabetic nephropathy codes for the numerator of the eQCM “Diabetes: Medical Attention for Nephropathy” were used; and for persistent asthma codes for the denominator of the eQCM “Use of appropriate medications for asthma” were used. An individual was treated as having one of these three conditions in 2016 if the EHR indicated a disease specific code on any date in 2016. Data were analyzed in 2018.

### Statistical analysis

Crude period prevalence (prevalent case count / study sample) during 2016 was calculated for each disease and each network, based on different study sample definitions that required 2,3,4,5 or 6 visits to the health care system between January 1 2016 and December 31 2016). This analysis was also repeated restricting the study population to patients between the ages of 46 and 65 in 2016. This project was approved by the CAPriCORN IRB.

## Results

Figure 1 shows how the study sample for each health system changed as the required number of visits to that health system during 2016 increased. In all cases, not surprisingly, increasing the minimum number of visits required dramatically reduced the size of the sample. Interestingly, the sites showed fairly similar percentage declines in the size of the sample as the number of required visits increased. For all three sites, requiring 6 visits nearly halved the sample size, compared to requiring only 2 visits.

**Figure 1.**
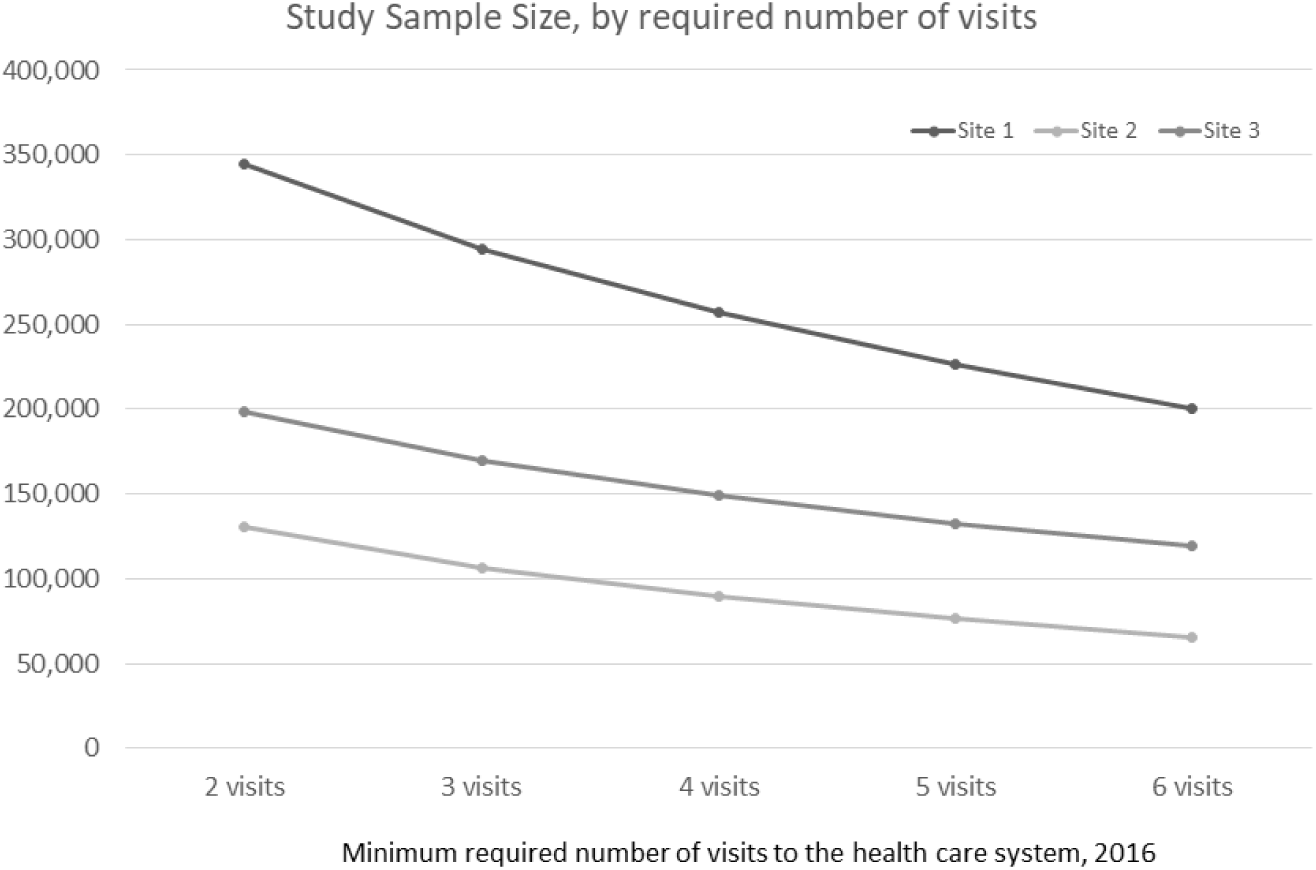
Study sample for each health system for different minimum required number of visits ([inpatient, outpatient, or emergency room visits] from January 1 2016 to December 31 2016)

Figure 2 shows how the calculations of crude period prevalence (2016) for MI (a) diabetic nephropathy (b) and persistent asthma (c) changed across health systems, as the minimum number of health system visits required for inclusion in the study sample increased. The number of patients with each of these diagnoses also fell as the minimum number of visits increased, but much more slowly than the denominator. As a result, the prevalence of all three diseases increased as the number of visits required to enter the study sample increased across all health systems. However, the rate of increase differed across sites and diseases. For site 2 increasing the number of required visits from 2 to 6 increased MI prevalence by 36% and increased persistent asthma prevalence by 48%. For MI, increasing the number of required visits from 2 to 6 increased MI prevalence at site 1 by 57% and MI prevalence at site 2 by 36%. To see if some standardization of populations across the three sites reduced observed differences in the way prevalence changed with increasing number of required visits, we also calculated crude period prevalence (2016) for MI, diabetic nephropathy, and persistent asthma by the minimum number of health system visits required for inclusion in the subset of 46-65 year olds (Supplementary Table 1). This attempt at standardization actually increased observed differences in the way MI prevalence changed with increasing number of required visits across sites; increasing the number of required visits from 2 to 6 in the subset of 46-65 increased MI prevalence at site 1 by 68% and MI prevalence at site 2 by 37%.

**Figure 2.**
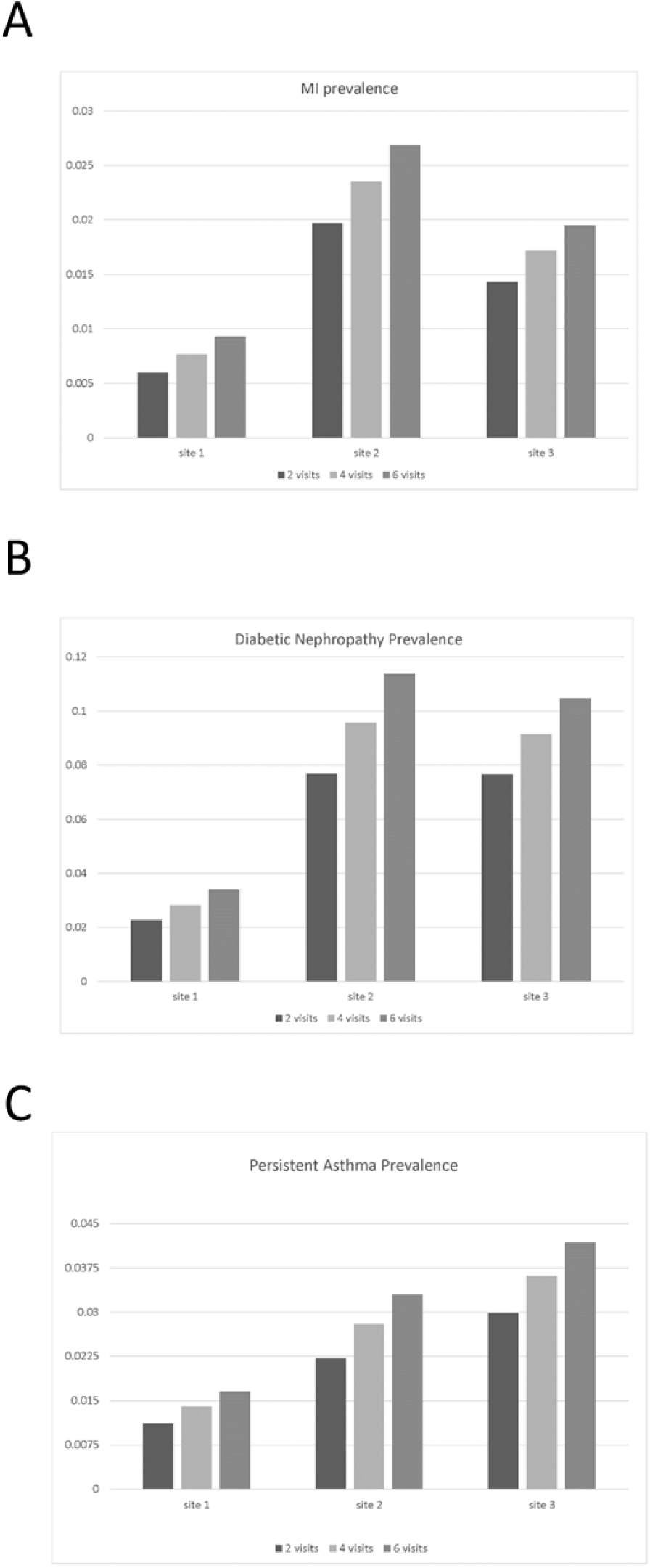
Variation in basic descriptive epidemiology metrics (example: period prevalence) with increase in minimum number of health care visits required to enter the study sample. Estimates of period prevalence during 2016 for (a) MI (b) diabetic nephropathy and (c) persistent asthma, for different minimum numbers of required visits during 2016.

## Discussion

This study demonstrates that for three disease conditions, across three different health systems, estimates of basic descriptive epidemiology metrics, like crude disease period prevalence, change as one increases the minimum number of health care system visits required to be included in the study sample. The increase in prevalence with minimum number of visits is not surprising; people with more contact with the health care system will have more complete records of their health status. However, also as one increases the number of visits to a health care system required to enter a study sample, one loses healthy individuals (who presumably have less contact with any health care system), making results less generalizable to the population. Unfortunately, there appears to be only limited consistency in the magnitude of increases in period prevalence estimates as one increases the minimum number of required visits, both across different diseases (within a single health care system) and across health care systems (for a single disease), which severely limits broadly generalizable recommendations about study sample definition using EHRs from non-closed systems. Given these challenges (as well as lack of adjustment, different measurement methods, and geographic and demographic diversity) it is not surprising that the calculated crude estimates of MI, diabetic nephropathy, and asthma differed both widely across health systems and from recent Behavioral Risk Factor Surveillance System and National Health Interview Survey estimates ^8–10^.

Researchers are developing methods to estimate disease prevalence using EHR data, some with notable success ^11–13^. This study suggests decisions in the basic definition of the study population can greatly impact estimation of disease prevalence, which, in turn, can greatly impact many descriptive and analytic epidemiology analyses. Study population definition should be a carefully considered element in any clinical or epidemiologic study using EHR data.

Investigators should carefully define the study sample (and their reasons for choosing that study sample definition, including information about the effect of choosing different definitions) when using EHRs in research. Investigators should also consider that different EHR study sample definitions may be required for the study of different diseases. If an algorithm for EHR study sample selection for a disease is proposed, it should be tested and validated across multiple health systems, in the same manner that EHR phenotype definitions are often validated ^14^.

This work was funded by CDRN-1306-04737-IC, CDRN-1306-04631, CDC-5U18DP006120-04-00, CDC 1U18DP006120-01, NCATS NIH UL1TR001436, NCATS NIH UL1TR002537, and NIDDK 5U18DP006120-02.

## Acknowledgments

The authors would like to acknowledge Zahra Hosseinian and Charon Gladfelter for all their administrative help with this project.

