## Supplementary Table 1 for "The effect of number of healthcare visits on study sample selection and prevalence estimates in electronic health record data"

**Supplementary Table 1.** Variation in period prevalence (2016) of three diseases with increase in minimum number of health care visits required to enter the study sample, in participants aged 46-65, by site.

| Disease | # of visits |  | Period Prevalence | | |
| --- | --- | --- | --- | --- | --- |
|  |  | Site1 | | Site 2 | Site 3 |
| Myocardial Infarction | 2 visits | 0.0013 | | 0.0030 | 0.0032 |
| Myocardial Infarction | 4 visits | 0.0017 | | 0.0040 | 0.0038 |
| Myocardial Infarction | 6 visits | 0.0022 | | 0.0048 | 0.0044 |
| Diabetic Nephropathy | 2 visits | 0.0103 | | 0.0320 | 0.0348 |
| Diabetic Nephropathy | 4 visits | 0.0132 | | 0.0415 | 0.0422 |
| Diabetic Nephropathy | 6 visits | 0.0161 | | 0.0510 | 0.0492 |
| Persistent Asthma | 2 visits | 0.0098 | | 0.0197 | 0.0234 |
| Persistent Asthma | 4 visits | 0.0125 | | 0.0248 | 0.0287 |
| Persistent Asthma | 6 visits | 0.0148 | | 0.0294 | 0.0335 |
